# Comparative genome-wide CRISPR-Cas9 screening reveals host factors for MHC-II-mediated influenza A virus infection

**DOI:** 10.64898/2026.11.25.767822

**Authors:** Laura M. Arroyo-Fernández, Lars Pache, Umut Karakus, Irina Glas, Patricia Martínez-Barragán, Silke Stertz

## Abstract

Virus-host interactions during influenza A virus (IAV) infection are key determinants of interspecies transmission and disease progression. The identification of IAVs capable of using sialic acid and major histocompatibility complex class II (MHC-II) as entry receptors raises the question of whether alternative receptor usage is associated with distinct host factor requirements. Here, we conducted comparative genome-wide CRISPR-Cas9 loss-of-function screens in human lung epithelial cells displaying either sialic acid or MHC-II, using a dual receptor-specific H2N2 strain. Our approach identified known and novel host genes required for sialic acid- and MHC-II-mediated IAV infection and, by integrating both receptor-associated datasets, revealed host factors specific for the MHC-II-mediated infection route. Among these, we uncovered syntaxin-2, a SNARE protein involved in vesicle trafficking, as a host factor specifically required for MHC-II-mediated IAV entry. We additionally identified multiple components of the type I interferon signaling pathway and show that virus-induced interferon signaling influences cell proliferation during survival-based screening. This study systematically characterizes host factor requirements for IAV infection via sialic acid and MHC-II and suggests that alternative receptor usage is associated with distinct cellular dependencies.

## INTRODUCTION

Influenza A viruses (IAVs) are respiratory pathogens that cause seasonal epidemics and occasional pandemics with substantial global health and economic impact. Seasonal influenza is estimated to result in 3-5 million cases of severe disease and up to 650,000 deaths annually (1, 2). In addition to circulating in humans, IAVs are maintained in diverse animal reservoirs, including birds and swine, from which zoonotic spillovers can occur and in rare instances give rise to novel pandemic strains with increased transmissibility and pathogenicity (3, 4).

Virus-host interactions during the initial stages of the viral infection cycle are major determinants of tropism and critically influence transmission and pathogenesis (5). Entry of conventional IAV strains is initiated by binding of the viral hemagglutinin (HA) to sialic acid-containing glycoconjugates at the cell surface (6, 7), followed by endocytic uptake and pH-dependent membrane fusion (8). The cellular mechanisms and host factors mediating sialic acid-dependent entry have been extensively characterized, advancing our understanding of IAV interspecies adaptation and providing novel targets for antiviral therapeutics (9–12).

Major histocompatibility complex class II (MHC-II) molecules have been recently identified as novel IAV entry receptors. MHC-II molecules are the unique receptors of the bat-derived H17N10 and H18N11 (13, 14) and avian H19 viruses (15). Moreover, they are capable of mediating sialic acid-independent entry of selected human and avian H2N2 (16) and H3N2 (17) strains. Because receptor engagement initiates the infection cycle, it is expected to play a critical role in determining which cellular mechanisms the virus exploits for infection. Previous studies have provided evidence for a direct interaction between HA and MHC-II (18) and suggested that, following MHC-II engagement, endosomal acidification is required to trigger HA-mediated membrane fusion and viral entry (19, 20). However, the cellular mechanisms involved in MHC-II-mediated IAV infection are poorly understood. Therefore, a comprehensive characterization of the host requirements for MHC-II receptor usage is needed to provide insights into its potential contribution to interspecies adaptation and identify novel targets for host-directed antiviral intervention.

Here, we established a receptor-defined cellular model to independently characterize the sialic acid- and MHC-II-mediated IAV infection routes. Using this system, we performed comparative genome-wide CRISPR-Cas9 loss-of-function screens based on cell survival following infection with a dual receptor-specific human H2N2 IAV to profile the host factor landscape associated with sialic acid- and MHC-II-mediated infection. Integration of both receptor-specific datasets enabled the prioritization of host factors selectively associated with MHC-II-mediated infection and identified syntaxin-2 (STX2), a SNARE protein involved in intracellular vesicle fusion, as a factor specifically required for efficient H2N2 IAV entry downstream of MHC-II receptor engagement. Overall, our study defines receptor-associated host factor requirements for IAV infection, enabling the characterization of the cellular pathways involved in MHC-II-mediated IAV entry.

## RESULTS

### A receptor-specific cellular model to dissect sialic acid- and MHC-II-mediated IAV entry

As human H2N2 IAVs can use both sialic acid and MHC-II for cell entry, they provide a suitable model to investigate and compare the host factors required for receptor-specific infection. To investigate sialic acid- and MHC-II-mediated entry pathways independently, we established a cellular model based on the respiratory epithelium-derived Calu-3 cell line (Fig. 1a). This system builds on a previously described double knockout (DKO) Calu-3 cell line (16), which lacks *SLC35A1*, encoding the CMP-sialic acid transporter, and *HLA-DRA*, encoding the alpha chain of the HLA-DR complex. From this DKO background, we selectively reconstituted receptor expression by lentiviral transduction of either *SLC35A1*, to restore sialic acid biosynthesis (DKO-SLC35A1), or *HLA-DRA* together with *HLA-DRB1*, to form functional HLA-DR complexes (DKO-HLA-DR). A cell line transduced with an empty lentiviral vector (DKO-Ctrl) was generated as a negative control. To assess receptor expression in the engineered cell lines, we quantified the surface levels of sialic acid and HLA-DR by flow cytometry. DKO-SLC35A1 cells displayed high levels of surface α2,3- and α2,6-linked sialic acid but lacked HLA-DR expression, whereas DKO-HLA-DR cells expressed HLA-DR but were deficient in sialic acid (Fig. 1b). Next, we tested if these cell lines support receptor-specific entry of IAVs. To do so, we used a panel of HIV-based luciferase-encoding virus-like particles (VLPs) pseudotyped with HAs and NAs from IAV strains with defined receptor specificity: the H1N1 strain A/Brisbane/59/2007 and the H3N2 strain A/Hong Kong/2286/2017 (sialic acid-dependent), the H18N11 strain A/bat/Peru/033/2010 (MHC-II-dependent), and the H2N2 strains A/Japan/305/1957 and A/Taiwan/1/1964 (dual specificity for sialic acid and MHC-II). Both the parental Calu-3 and the receptor-specific cell lines supported HA-mediated VLP entry, while the corresponding control cells were resistant to infection (Fig. 1c). As expected, H2-VLPs entered cells displaying either sialic acid or HLA-DR. In contrast, H1- and H3-VLPs entered exclusively sialic acid-positive cells and bat-derived H18-VLPs specifically entered cells expressing HLA-DR. To extend these observations to replication-competent viruses, we performed a single-cycle infection assay using authentic H1N1 (A/Puerto Rico/8/1934) and H2N2 (A/Japan/305+/1957) viruses and measured viral protein expression at 7 h post-infection. Consistent with the previous assay, infection with the H1N1 strain A/Puerto Rico/8/1934 was only observed in sialic acid-positive cells (parental Calu-3 and DKO-SLC35A1), whereas the H2N2 strain A/Japan/305+/1957 infected both sialic acid- and HLA-DR-positive cells (parental Calu-3, DKO-SLC35A1 and DKO-HLA-DR) (Fig. 1d). Together, these results validate our cellular model for the independent assessment of receptor-specific IAV entry pathways.

**Figure 1.**
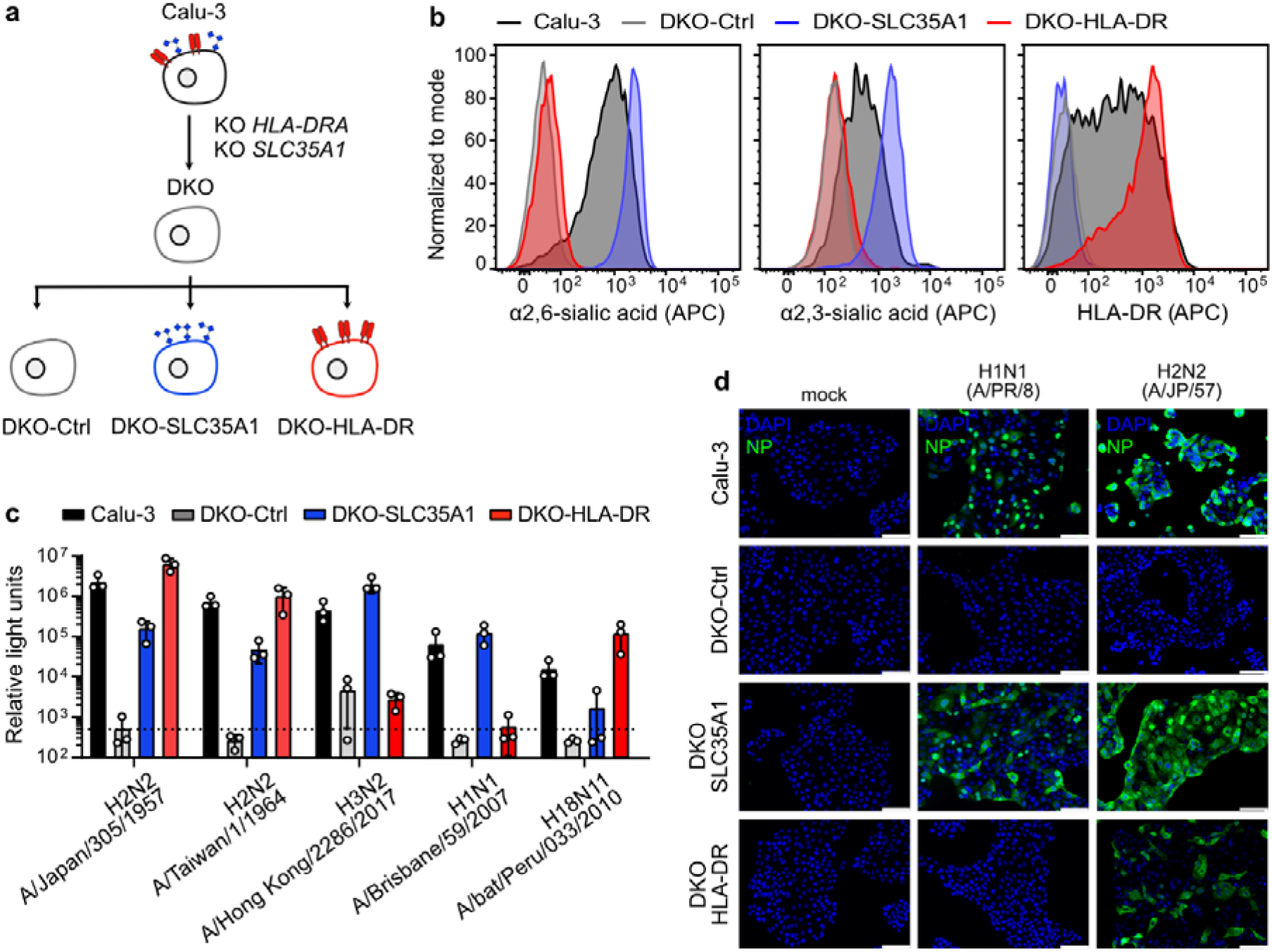
Calu-3 DKO receptor-specific cellular model. **(a)** Schematic representation of the cellular model. Calu-3 *HLA-DRA*/*SLC35A1* DKO cells were transduced with *SLC35A1* (DKO-SLC35A1), *HLA-DRA* and *HLA-DRB1* (DKO-HLA-DR) or an empty vector control (DKO-Ctrl) to generate receptor-specific cell lines. **(b)** Surface expression levels of HLA-DR, α2,3-, and α2,6-linked sialic acids were determined by staining with an APC-conjugated anti-HLA-DR antibody, biotinylated *Maackia amurensis* II lectin, and biotinylated *Sambucus nigra* lectin, respectively, and analyzed by flow cytometry. Representative histogram from n = 2 biological replicates. **(c)** Cells described in (a) were infected with HIV-based luciferase-encoding VLPs pseudotyped with glycoproteins (HA and NA) from the indicated IAV strains. Luciferase activity was measured at 72 h post-infection. Data are means ± s.d. from n = 3 biological replicates. Dashed line indicates limit of detection. **(d)** Cells described in (a) were infected with A/Puerto Rico/8/34 or A/Japan/305+/1957 at MOI 5 and fixed at 7 h post-infection. Viral nucleoprotein (NP; green) and nuclei (DAPI; blue) were detected by immunofluorescence. Scale bar = 100 µm. Image representative of n = 2 biological replicates.

### Comparative CRISPR-Cas9 screens identify host factors specific for sialic acid- or MHC-II-mediated H2N2 IAV infection

To systematically identify host factors required for receptor-dependent infection, we performed comparative genome-wide CRISPR-Cas9 loss-of-function screens (Fig. 2a). DKO-SLC35A1 and DKO-HLA-DR cells were transduced with the human GeCKOv2 sgRNA library (21) to generate pooled knockout (KO) populations that were subsequently infected with the cytopathic H2N2 IAV strain A/Japan/305+/1957. Prior to screening, we assessed the susceptibility of receptor-specific cell lines to virus-induced cytopathic selection. Infection with H2N2 IAV induced robust cell death in both DKO-SLC35A1 and DKO-HLA-DR cells, while DKO-Ctrl cells remained resistant (Fig. 2b), confirming that virus-induced cell death is dependent on receptor expression and providing a suitable dynamic range for selection. After infection, genomic DNA from surviving cells was harvested and sgRNA abundances in pre-and post-selection populations were quantified by deep sequencing (Fig. S1a, Supp. Table S1). Screen data was analyzed using the PinAPL-Py bioinformatics tool (22) and genes were ranked based on the adjusted robust rank aggregation (αRRA) algorithm (Supp. Tables S2-S3). Screens were performed in biological duplicates.

**Figure 2.**
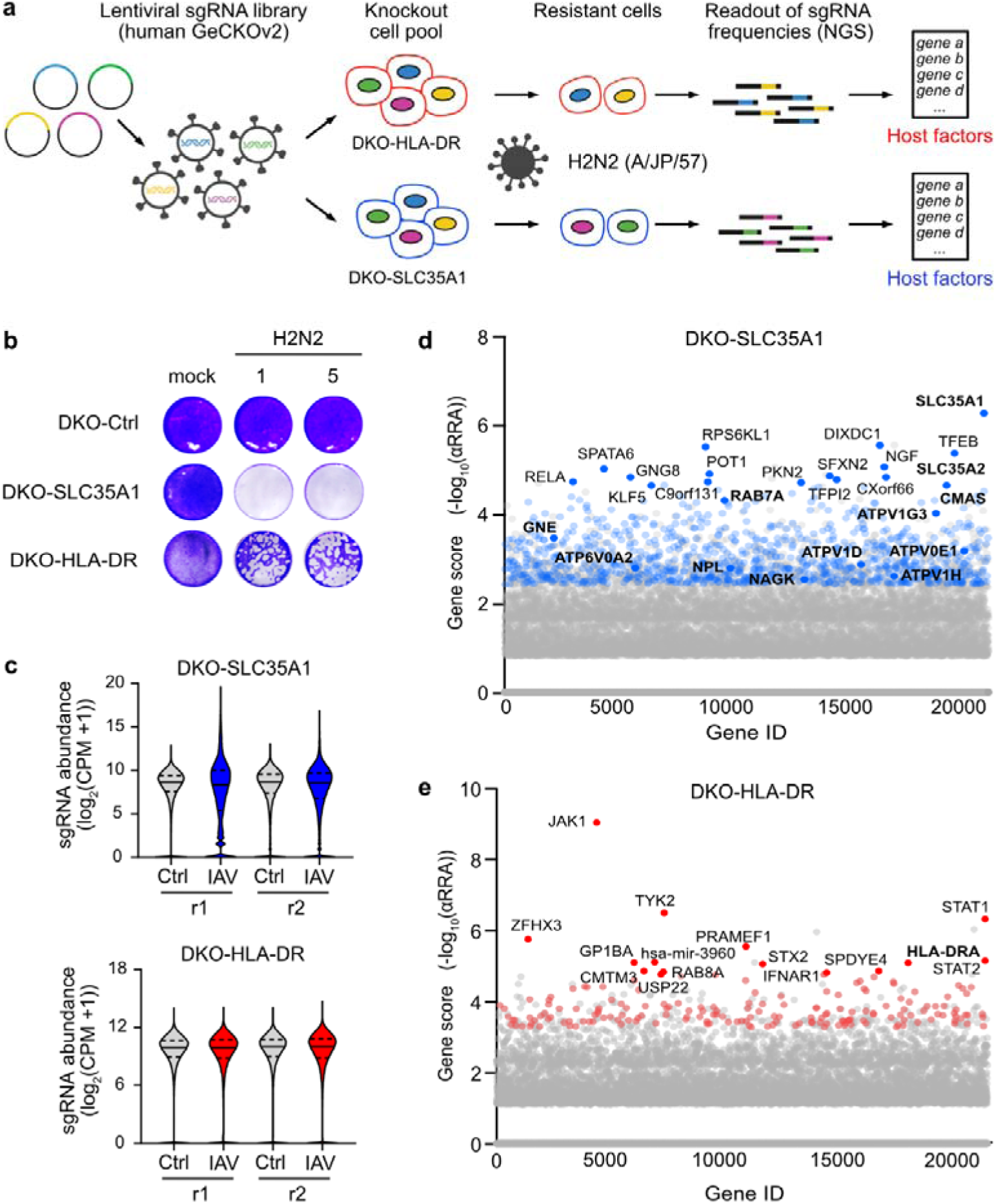
Comparative CRISPR-Cas9 screens identify host factors specific for sialic acid- or MHC-II-mediated H2N2 IAV infection. **(a)** Schematic of the comparative CRISPR-Cas9 screening strategy to identify host factors required for either sialic acid- or HLA-DR-mediated H2N2 IAV infection. **(b)** Cells were infected with A/Japan/305+/1957 (MOI 1 or 5). At 72 h post-infection, cells were fixed and stained with crystal violet to visualize virus-induced cytopathic effect. Representative image from n = 2 independent experiments. **(c)** Distribution of sgRNA counts in DKO-SLC35A1 (grey/blue) or DKO-HLA-DR (grey/red) cells before (Ctrl) and after (IAV) infection with A/Japan/305+/1957. Read counts were averaged across two biological replicates, normalized using size-factor normalization, and log_2_-transformed with a pseudocount of 1 (log_2_(CPM + 1)). Distributions are visualized as violin plots, with median (solid line) and interquartile range (dashed line) indicated. **(d, e)** Gene scores from CRISPR-Cas9 screens in DKO-SLC35A1 (d) and DKO-HLA-DR (e). Scores were calculated using the αRRA algorithm implemented in the PinAPL-Py tool. Significant hits (p <0.01 and ≥3 significant sgRNAs per gene) are highlighted in color. The top 15 genes and a selected subset of previously described host factors (bold) are annotated.

Analysis of sgRNA abundance revealed a marked shift in library composition following infection in DKO-SLC35A1 cells, indicative of a strong selective pressure (Fig. 2c). Initial screening in DKO-HLA-DR cells at multiplicity of infection (MOI) 1 resulted in a more modest shift in the sgRNA distribution, consistent with a reduced selective pressure. To enhance selection, a second screen replicate was performed using two rounds of infection at MOI 5, which yielded a more pronounced enrichment of sgRNAs. The observed changes in sgRNA distribution are quantitatively reflected in the corresponding Gini coefficients of pre- and post-selected populations (Fig. S1b). Across both screens, high reproducibility between biological replicates was observed, as indicated by Pearson correlation analysis (Fig. S1c).

Applying significance thresholds (p-value <0.01 and ≥3 enriched sgRNAs per gene), the screen in DKO-SLC35A1 cells identified 1,083 candidate genes (Fig. 2d). Notably, we recovered several established IAV host factors (9, 11, 23), including multiple components of the sialylation pathway (*SLC35A1*, *SLC35A2*, *CMAS*, *GNE*, *NAGK*, and *NPL*) and subunits of the vacuolar ATPase (V-ATPase) complex (*ATP6V1H*, *ATP6V1D*, *ATP6V1G3*, *ATP6V0A2*, and *ATP6V0E1*). In DKO-HLA-DR cells, 186 genes were significantly enriched, with *HLA-DRA* ranking among the top hits (Fig. 2e). Differences in the number of significant hits identified between screens likely reflect the different selection strengths observed under the two receptor-specific conditions. Despite this difference, both datasets recovered established IAV host factors, indicating that our comparative screening strategy effectively captures biologically relevant proviral host factors.

### Pathway analysis and integration of CRISPR-Cas9 screens reveal MHC-II-specific host factors

To characterize the cellular pathways represented in each screen, we performed Gene Ontology (GO) enrichment analysis of the hits identified in the sialic acid and HLA-DR screens using Enrichr (24–26) (Fig. 3a, Supp. Table S4). In the sialic acid screen, enriched biological processes included multiple metabolic pathways, such as carboxylic acid, sulfur amino acid and N-acetylneuraminic acid metabolism, the latter reflecting the dependency of viral entry on sialic acid in DKO-SLC35A1 cells. Pathways associated with endosomal trafficking, lysosomal organization and intracellular pH regulation were also identified, consistent with the internalization of IAV through receptor-mediated endocytosis (27). In addition, processes related to cell adhesion, migration and epithelial remodeling, including lamellipodium assembly, were also found, in line with previous evidence implicating cytoskeletal and membrane remodeling processes in IAV entry (28). In contrast, the HLA-DR screen showed a prominent enrichment of interferon- and cytokine-mediated signaling pathways, including JAK-STAT signaling and cellular responses to type I interferon (IFN-I), likely related to the activation of innate antiviral responses following IAV infection. Pathways associated with oxidative stress, cytochrome c release, apoptosis and regulation of necroptosis were also enriched, suggesting a role for cellular stress and cell-death pathways in survival following infection. Processes related to ceramide, sphingolipid and lipid biosynthesis, as well as endoplasmic reticulum protein localization and retention, were also represented. Overall, the two screens displayed distinct patterns of pathway enrichment. While the sialic acid screen predominantly recovered processes associated with conventional IAV entry, the HLA-DR screen was characterized by a stronger representation of pathways associated with cellular responses to infection. These observations suggest that engagement of sialic acid and MHC-II receptors may differentially influence cellular pathways and responses during IAV infection and motivate a more in-depth characterization of the host requirements underlying receptor-specific infection.

**Figure 3.**
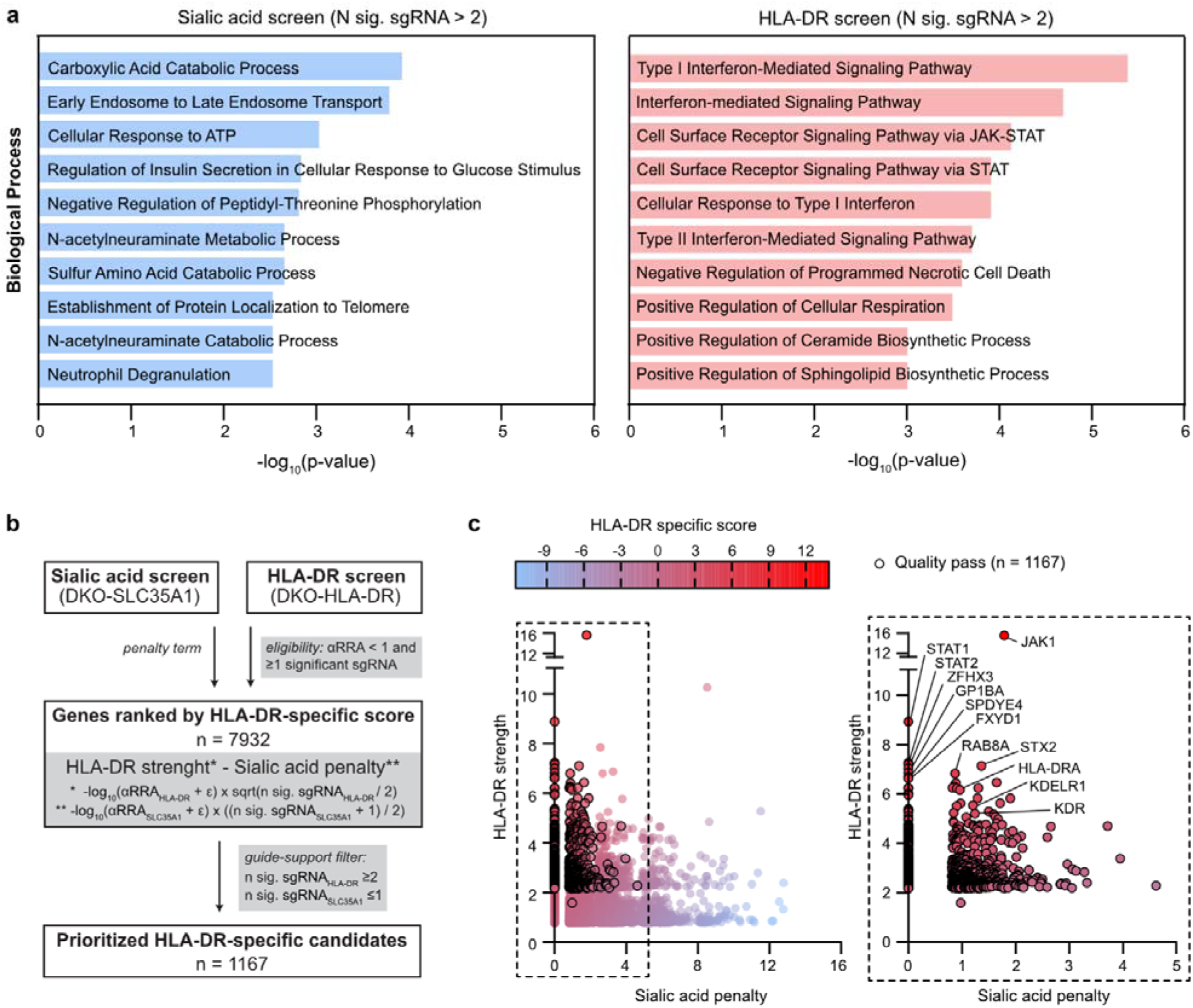
Pathway analysis and integration of CRISPR-Cas9 screens reveal MHC-II-specific host factors. **(a)** Gene ontology analysis of hits (p-value <0.01 and ≥3 enriched sgRNAs per gene) identified in the screens performed in DKO-SLC35A1 (blue) and DKO-HLA-DR (red) cells. **(b)** Schematic overview of the integrated scoring strategy used to prioritize HLA-DR-specific host factors. The analysis starts with genes identified in the HLA-DR screen, calculates an HLA-DR-specific score by integrating results from both receptor-specific screens, and applies guide-support filters to define a prioritized set of HLA-DR-specific candidates. **(c)** Scatter plot showing the HLA-DR strength and sialic acid penalty for each gene identified in the DKO-HLA-DR screen. Genes are colored according to their HLA-DR-specific score. Genes passing the guide-support criteria (≥2 significantly enriched sgRNAs in the HLA-DR screen and ≤1 significantly enriched sgRNA in the sialic acid screen) are outlined. The right panel shows an enlarged view of the top-ranked region, highlighting genes that pass the guide-support filter and labeling selected candidate genes.

Next, we targeted our analysis to the identification of host factors selectively associated with HLA-DR-mediated IAV infection. Because DKO-HLA-DR and DKO-SLC35A1 cells share a common DKO background but differ in receptor availability for viral entry, each screen is expected to recover receptor-specific factors as well as host factors shared between entry routes. To distinguish HLA-DR-specific candidates from shared IAV host factors, we integrated the two screen datasets using an HLA-DR-specific scoring strategy (Fig. 3b). This approach combines gene-level enrichment significance (αRRA) with the number of significantly enriched sgRNAs. It prioritizes genes with stronger support in the HLA-DR screen while penalizing genes also enriched in the sialic acid screen. A detailed description of the scoring strategy is provided in the Methods section. Application of this approach generated a ranked list of candidate HLA-DR-specific host factors (Fig. 3c, Supp. Table S5). To define a prioritized candidate subset, we required each gene to be supported by at least two significantly enriched sgRNAs in the HLA-DR screen and no more than one significantly enriched sgRNA in the sialic acid screen. Applying these criteria to the ranked list identified 1,167 prioritized candidates (Fig. 3c). *HLA-DRA* ranked among the highest-scoring candidates (rank 21), demonstrating that the ranking strategy recovered a known receptor-specific factor.

### The IFN-I response limits cell growth during H2N2 IAV infection

IFN-I signaling emerged as a prominent feature associated with MHC-II-mediated infection in our comparative screening. Pathway analysis of candidate genes identified in the HLA-DR screen revealed significant enrichment of multiple IFN-related pathways. At the gene level, *IFNAR1*, *IFNAR2*, *JAK1*, *TYK2*, *STAT1*, *STAT2*, and *IRF9* were among the top-enriched hits, with *JAK1*, *STAT1*, *STAT2*, and *IRF9* also ranking among the top hits in the HLA-DR-specific analysis. This result was counterintuitive given the established antiviral role of IFN-I signaling in IAV infection, so we next investigated the contribution of IFN-I signaling in HLA-DR-mediated IAV infection.

To do so, we selected representative genes involved in the IFN-I signaling cascade - *IFNAR2* and *STAT1* and generated polyclonal KO populations in DKO-HLA-DR cells using CRISPR-Cas9. Cells transduced with a non-targeting sgRNA (sgNT) or a sgRNA targeting *HLA-DRA* were included as negative and positive controls, respectively. Efficient depletion of IFNAR2, STAT1 or the HLA-DR complex was confirmed by flow cytometry (Fig. 4a-c). We first assessed whether these factors are required for single cycle H2N2 IAV infection. Disruption of *IFNAR2* or *STAT1* did not reduce viral nucleoprotein (NP) expression at 7 h post-infection compared to control cells (Fig. 4d). In contrast, KO of *HLA-DRA* markedly impaired NP expression, consistent with its role as a receptor in this cell line. These results indicate that IFN-I signaling components are dispensable for viral entry and viral gene expression. We next evaluated their role in multicycle IAV replication. Viral growth kinetics were evaluated using a recombinant H2N2 JP/57 6+2 virus, which encodes the surface HA and NA proteins of A/Japan/305/1957 and the internal segments of A/Puerto Rico/8/1934. This virus retains the dual receptor specificity of the wild-type H2N2 virus (Fig. S2) but displayed an enhanced replication capacity in cell culture. Over multiple days post-infection, growth of recombinant JP/57 6+2 virus was comparable between control and IFNAR2-or STAT1-deficient cells, whereas depletion of HLA-DR resulted in a pronounced reduction in viral titers (Fig. 4e). These findings demonstrate that IFN-I signaling factors do not function as proviral host factors for IAV infection in this model.

**Figure 4.**
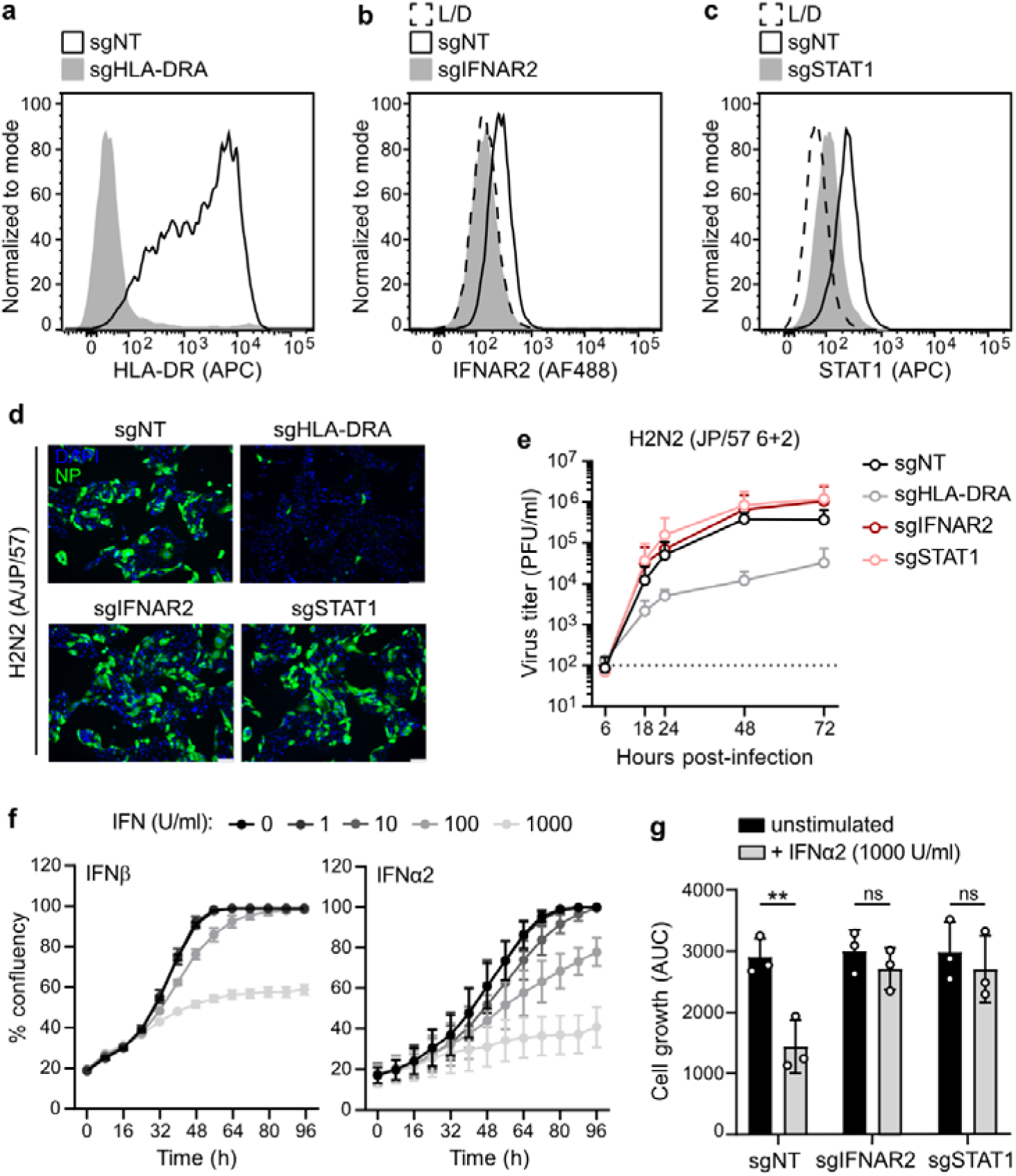
The IFN-I response limits cell growth during IAV infection. DKO-HLA-DR cells were transduced with sgRNAs targeting indicated genes or a non-targeting control (sgNT) and selected for 7 days. **(a-c)** Surface expression of HLA-DR (a), IFNAR2 (b) and intracellular expression of STAT1 (c) were measured by flow cytometry. Histograms representative from n = 2 independent experiments. **(d)** Cells described in (a-c) were infected with A/Japan/305+/1957 at MOI 5. At 7 h post-infection cells were fixed and viral nucleoprotein (NP; green) and nuclei (DAPI; blue) were detected through immunofluorescence. Scale bar = 100 µm. Images representative of n = 2 independent experiments. **(e)** Cells were infected with H2N2 JP/57 6+2 at MOI 0.5 and viral titers (PFU/mL) were determined by plaque assay at the indicated time points. Dashed line indicates limit of detection. Data are means ± s.d. from n = 3 independent experiments. **(f)** DKO-HLA-DR cells transduced with sgNT were seeded at approximately 20% confluency and stimulated with the indicated concentrations of IFNβ or IFNα2. Cell confluency was measured every 8 h for 96 h using an Incucyte S3. The percentage of confluency over time is represented. Data are means ± s.d. from n = 3 independent experiments. **(g)** The indicated sgRNA-transduced cell lines were seeded at approximately 20% confluency and stimulated with 1000 U/mL IFNα2. Cell confluency was monitored every 8 h for 96 h using an Incucyte S3. The area under the curve (AUC) of the cell confluency versus time curves is represented. Data are means ± s.d. from n = 3 independent experiments. Statistical significance was determined using two-tailed unpaired t-test. ns, not significant; \**\*P* ≤ 0.01.

Given that our screening strategy relies on cell survival following virus infection, we reasoned that host factors influencing cell fitness under infection conditions could also be selected. Given that IFN-I is known to exert cytostatic effects in addition to its antiviral activity (29), we next examined the impact of exogenous IFN-I stimulation on cell proliferation. Treatment of DKO-HLA-DR sgNT cells with increasing concentrations of either IFNβ or IFNα2 resulted in a dose-dependent inhibition of cell growth (Fig. 4f), indicating that IFN-I signaling limits proliferation in this system. To determine if this effect is mediated through canonical IFN-I signaling, we compared the cell growth of control and KO cells after IFN-I stimulation. While exposure to IFNα2 significantly reduced proliferation of control cells, IFNAR2- and STAT1-deficient cells were largely resistant to IFN-I-induced growth inhibition (Fig. 4g). Together, these results suggest that the enrichment of IFN-I signaling factors is related to the influence of antiviral signaling on cell proliferation, rather than a direct requirement for IAV replication.

### Validation of HLA-DR-specific host factors

To validate HLA-DR-specific entry factors for IAV, we selected a subset of top-ranked hits based on their potential relevance to viral entry processes. We prioritized candidates involved in membrane organization and trafficking, including both surface-exposed receptors (*KDR*) and intracellular trafficking regulators (*STX2*, *KDELR1*). These factors are expected to influence key steps for IAV entry, such as viral attachment, endocytosis, and membrane fusion. We first determined whether these factors contribute to HLA-DR-mediated infection. For each gene, DKO-HLA-DR cells were transduced with a pool of the three top sgRNAs enriched in the screen to generate polyclonal KO cell populations. Cells transduced with sgNT and a sgHLA-DRA were included as negative and positive controls, respectively. KO cells were subsequently infected with the human H2N2 strain A/Japan/305+/1957, and synthesis of the viral M segment was quantified by digital PCR at 7 h post-infection. As expected, disruption of *HLA-DRA* markedly reduced viral infection. Among the selected candidates, genetic depletion of *STX2* and *KDR* consistently decreased viral M segment production, while depletion of *KDELR1* produced a more variable phenotype (Fig. 5a). To facilitate subsequent validation experiments, we next employed a single-cycle H2N2 (A/JP/57)-*Renilla* reporter virus. Consistent with the results obtained using authentic virus, depletion of *STX2* and *KDR* significantly reduced reporter virus infection, while the effect was more modest for *KDELR1* (Fig. 5b).

**Figure 5.**
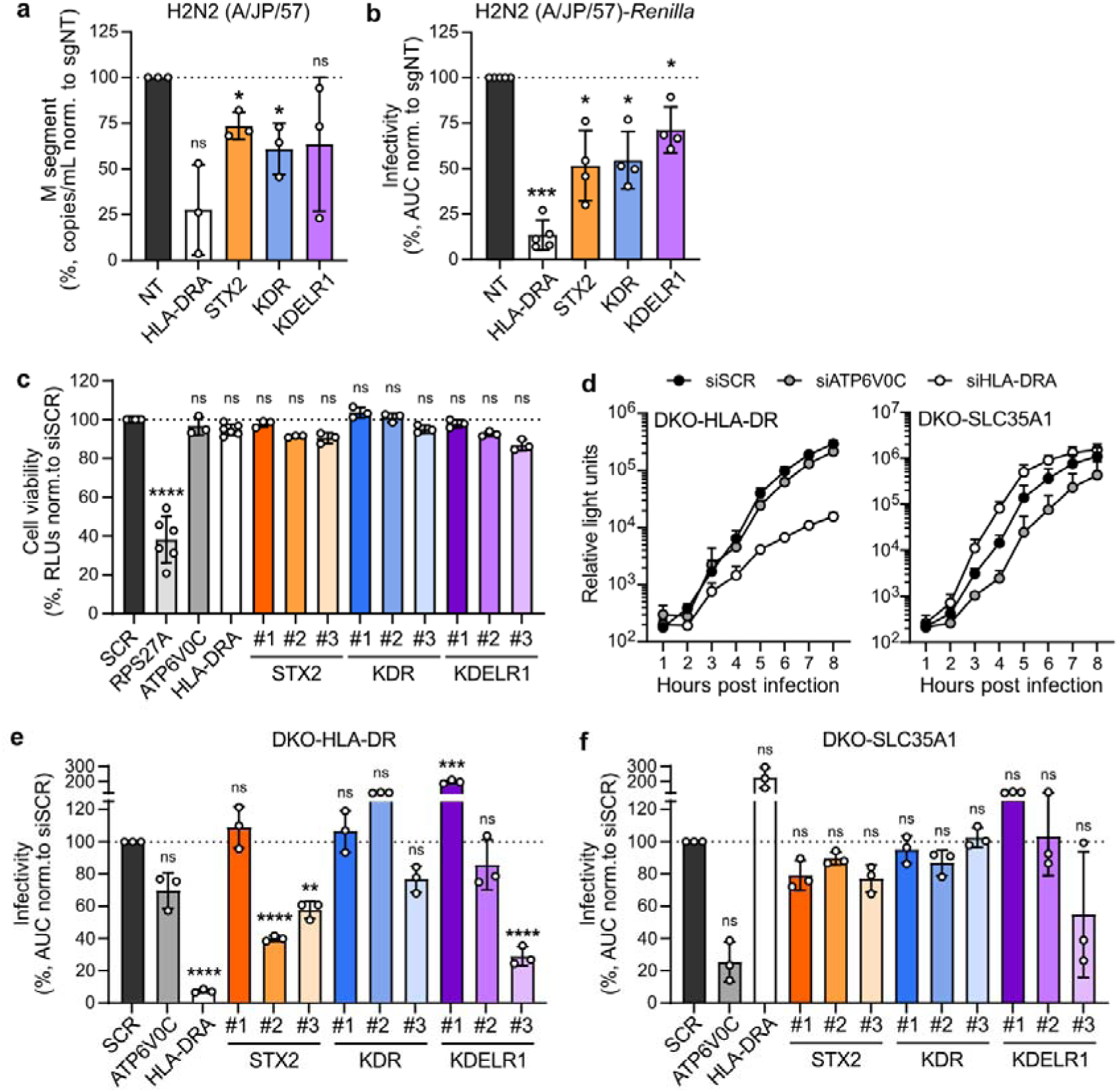
Validation of HLA-DR-specific host factors. **(a)** DKO-HLA-DR cells transduced with sgRNAs targeting indicated genes or a non-targeting control (sgNT) were infected with A/Japan/305+/1957 (MOI 0.5). Viral M segment copy numbers were quantified by digital PCR at 7 h post-infection and normalized to the sgNT control. Dashed line at 100%. Data are means ± s.d. from n = 3 independent experiments. **(b)** DKO-HLA-DR cells transduced with indicated sgRNAs were infected with H2N2 (A/JP/57)-*Renilla* (MOI 1). *Renilla* luciferase activity was measured for 8 h as a proxy of virus replication. The area under the curve (AUC) normalized to the sgNT control is shown. Dashed line at 100%. Data are means ± s.d. from n = 4-5 independent experiments. **(c)** DKO-HLA-DR cells were transfected with the indicated siRNAs for 48 h. Cell viability was quantified using CellTiter Glo. Data are means ± s.d. from n = 3 independent experiments. **(d-f)** DKO-HLA-DR and DKO-SLC35A1 cells were transfected with siRNAs targeting the indicated genes or a scrambled control (siSCR) and infected 48 h later with H2N2 (A/JP/57)-*Renilla* (MOI 1). **(d)** Representative *Renilla* luciferase kinetic curves. **(e, f)** AUC values of *Renilla* luciferase curves normalized to the siSCR control are shown. Dashed line at 100%. Data are means ± s.d. from n = 3 independent experiments. Statistical significance was determined in log_10_-transformed data using one-tailed paired t-test (a, b) or ordinary one-way analysis of variance (ANOVA) with Dunnett’s multiple comparisons test (c, e, f). ns, not significant; \**P* ≤ 0.05, \*\**P* ≤ 0.01, \*\*\**P* ≤ 0.001, \*\*\*\**P* ≤ 0.0001.

We next investigated whether these factors contribute specifically to HLA-DR-mediated infection or are also required for infection via sialic acid. To directly compare host factor requirements between the two receptor-specific cell lines, we performed siRNA-mediated gene knockdown. This strategy enables parallel gene silencing under identical experimental conditions while minimizing the variability associated with generating independent polyclonal KO cell populations. Each candidate gene was targeted by three independent siRNAs. In addition, siRNAs targeting *HLA-DRA* and *ATP6V0C*, a V-ATPase subunit known to be essential for sialic acid-dependent IAV entry (9), were included as receptor-specific controls. First, we confirmed that cell viability was not affected by any of the gene-targeting siRNAs tested (Fig. 5c). A siRNA targeting the essential gene *RPS27A* was used as a positive control for cell viability. Then, siRNA-transfected cells were infected with H2N2 (A/JP/57)-*Renilla* virus and infection was monitored over the first 8 h by measuring *Renilla* luciferase activity. Representative replication kinetics for the control conditions are shown in Fig. 5d, while quantification of the area under the curve for all conditions is shown in Fig. 5e, f. In DKO-HLA-DR cells, knockdown of *HLA-DRA* strongly impaired infection. Among the candidate genes, two of three siRNAs targeting *STX2* and one of three targeting *KDELR1* significantly reduced infection (Fig. 5e). However, knockdown of *KDR* did not significantly affect infection, although knockdown efficiency was not assessed. In DKO-SLC35A1 cells, knockdown of *ATP6V0C* markedly reduced infection, while knockdown of *HLA-DRA* had no inhibitory effect. In contrast to the phenotype observed in HLA-DR-expressing cells, knockdown of *STX2* or *KDELR1* produced only minor, non-significant effects in sialic acid-expressing cells (Fig. 5f).

Among the candidates tested, STX2 showed the strongest phenotype across the different genetic perturbation approaches, with reduced H2N2 infection observed following both CRISPR-Cas9-mediated KO and siRNA-mediated knockdown. Importantly, this effect was retained across two independent siRNAs and was not observed upon *STX2* knockdown in sialic acid-expressing cells, supporting a selective contribution to HLA-DR-mediated infection. Together, these data identify STX2 as a host factor that selectively promotes HLA-DR-mediated IAV infection and prompted us to further investigate its role during viral entry.

### STX2 is a host factor required for HLA-DR-mediated H2N2 IAV entry

STX2 is a Qa-SNARE protein involved in intracellular vesicle trafficking and membrane fusion (30, 31). Other members of the syntaxin family have previously been implicated in the entry of IAV and other negative-strand RNA viruses (32), raising the possibility that STX2 may similarly contribute to an early stage of infection. Given the selective effect of STX2 depletion on HLA-DR-expressing cells, we next investigated the contribution of STX2 to HLA-DR-mediated IAV entry.

First, we assessed knockdown efficiency of the three STX2-targeting siRNAs. At 48 h post-siRNA transfection, STX2 expression was reduced for all three siRNAs, with siSTX2#2 and siSTX2#3 producing a substantially stronger depletion (Fig. 6a, b). This correlated with the siRNA-specific reduction in H2N2 (A/JP/57)-*Renilla* infection previously observed. To validate these observations using authentic IAV, STX2-depleted cells were infected with the H2N2 A/Japan/305+/1957 strain, and viral M segment levels were quantified by digital PCR at 7 h post-infection. Consistent with the results obtained using the *Renilla*-reporter virus, siSTX2#2 and siSTX2#3 reduced viral RNA production of authentic H2N2 IAV (Fig. 6c).

**Figure 6.**
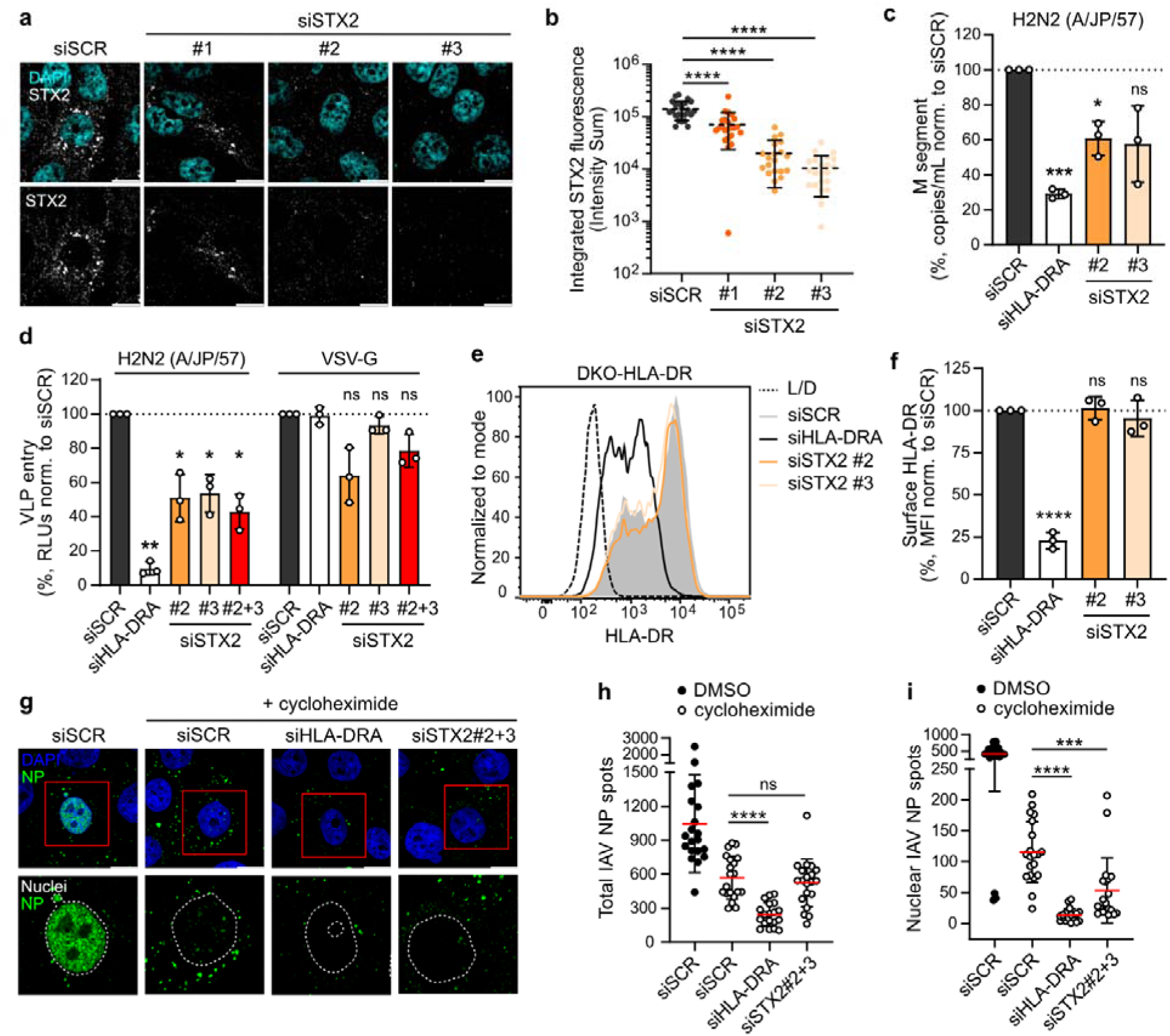
STX2 is a host factor required for HLA-DR-mediated IAV entry. DKO-HLA-DR cells were transfected with STX2-targeting siRNAs (siSTX2) or a scrambled control (siSCR) for 48 h. **(a)** siRNA-transfected cells were fixed, permeabilized, and stained for human STX2 (white) and nuclei (DAPI, cyan). Representative confocal images from n = 2 independent experiments are shown. Scale bar = 10 µm. **(b)** STX2-positive structures from samples described in (a) were identified by surface rendering using Imaris software, and the integrated fluorescence signal was quantified as the Intensity Sum of all STX2-positive surfaces per image. Each datapoint represents one confocal image from n = 2 independent experiments. Means ± s.d. is shown. **(c)** siRNA-transfected cells were infected with A/Japan/305+/1957 (MOI 0.5). Viral M segment copy numbers were quantified by digital PCR at 7 h post-infection and normalized to the sgNT control. Dashed line at 100%. Data are means ± s.d. from n = 3 independent experiments. **(d)** siRNA-transfected cells were infected with luciferase-encoding VSVΔG VLPs pseudotyped with HA and NA from the A/Japan/305/1957 (H2N2) strain or the glycoprotein from VSV (VSV-G). At 8 h post-infection luciferase signal was measured. Dashed line at 100%. Data are means ± s.d. from n = 3 independent experiments. **(e)** siRNA-transfected cells were fixed, stained and surface HLA-DR was quantified by flow cytometry. Representative histograms from n = 3 independent experiments. **(f)** Median fluorescence intensity (MFI) of samples described in (e) normalized to siSCR. Dashed line at 100%. Data are means ± s.d. from n = 3 independent experiments. **(g)** siRNA-transfected cells were infected with H2N2 JP/57 6+2 at MOI 40 in the presence of cycloheximide (100 µg/mL) or DMSO. At 4 h post-infection, cells were fixed, permeabilized and stained for viral NP (NP; green) and nuclei (DAPI; blue or dashed while line). Representative confocal images from n = 2 independent experiments are shown. Scale bar = 10 µm. **(h)** NP-positive puncta were identified and quantified by spot detection in Imaris. **(i)** For intranuclear NP quantification, nuclei were segmented based on the DAPI signal and NP puncta inside the nuclear surface were quantified. (h, i) Each datapoint represents one confocal image from n = 2 independent experiments. Means (red) ± s.d. is shown. Statistical significance was determined using two-tailed unpaired t-test (b, f, h, i), one-tailed paired t-test in log_10_-transformed data (c), two-tailed paired t-test in log_10_-transformed data (d). ns, not significant; \**P* ≤ 0.05, \*\**P* ≤ 0.01, \*\*\**P* ≤ 0.001, \*\*\*\**P* ≤ 0.0001.

We next asked if STX2 contributes specifically to viral entry. DKO-HLA-DR cells were infected with luciferase-encoding VSVΔG VLPs pseudotyped with either the HA and NA glycoproteins of H2N2 (A/Japan/305/1957) or the VSV glycoprotein (VSV-G). STX2 depletion selectively impaired infection by H2N2-pseudotyped particles while having little effect on VSV-G-mediated entry (Fig. 6d), indicating that STX2 specifically promotes HA-dependent entry into HLA-DR-expressing cells. Because STX2 has been primarily associated with recycling endocytosis and protein delivery to the plasma membrane (30, 33), we next examined if it modulates surface expression of the HLA-DR receptor. We quantified surface HLA-DR levels after siRNA-mediated knockdown of *STX2*. Flow cytometry analysis showed that knockdown of *STX2* did not alter HLA-DR surface levels (Fig. 6e, f), in contrast to the reduction observed after knockdown of *HLA-DRA*. These findings indicate that the observed entry defect is not a consequence of reduced receptor availability.

To further define the stage of IAV entry at which STX2 acts, we quantified incoming viral particles and assessed their intracellular localization, using nuclear accumulation of NP as a proxy for successful completion of the entry process. To specifically monitor incoming virions and exclude the contribution of newly synthesized viral proteins, cells were treated with cycloheximide, an inhibitor of *de novo* protein synthesis. At 48 h post-siRNA transfection, cells were infected with a high dose of H2N2 JP/57 6+2 in the presence of cycloheximide, and viral NP was detected by confocal microscopy at 4 h post-infection (Fig. 6g). As expected, cycloheximide treatment markedly reduced the NP signal compared with the DMSO-treated control (Fig. 6g, first two left panels), supporting the use of this approach to specifically monitor incoming viral particles. We subsequently quantified the total number of cell-associated NP-positive puncta as a measure of viral particle internalization (Fig. 6h), as well as the subset of NP-positive puncta localized within the nucleus as an indicator of successful progression through the entry pathway and nuclear delivery of viral ribonucleoproteins (vRNPs) (Fig. 6i). To validate nuclear NP accumulation as a readout of successful viral entry, we treated cells with bafilomycin A1, an inhibitor of endosomal acidification that blocks viral membrane fusion without preventing particle internalization (34, 35). Bafilomycin A1 treatment significantly reduced the number of nuclear NP puncta without affecting the total amount of cell-associated virions (Fig. S3), further supporting the use of nuclear NP accumulation to determine particles that have successfully completed the entry process. We next examined the effects of HLA-DR and STX2 depletion on viral particle internalization and nuclear delivery. *HLA-DRA* knockdown reduced both the total number of cell-associated NP puncta and the number of nuclear NP puncta, consistent with its established role in virion attachment. In contrast, STX2 depletion did not substantially affect the total number of cell-associated NP puncta but significantly reduced their nuclear accumulation. These findings indicate that, although viral particles can still attach in the absence of STX2, their subsequent progression through the entry pathway and delivery of vRNPs to the nucleus are impaired. Together, these findings confirm that STX2 contributes to HLA-DR-mediated IAV entry and support a role for STX2 at a post-attachment stage of the entry process.

## DISCUSSION

Receptor usage is a major determinant of IAV cellular and host tropism, shaping both cell susceptibility and the mechanisms underlying viral entry. While the molecular mechanisms governing sialic acid-mediated entry have been extensively characterized (36), much less is known about the cellular machinery that supports infection through the alternative receptor MHC-II. Here, we combine CRISPR-Cas9 screening with receptor-defined cellular models to characterize the host factor requirements for sialic acid- and HLA-DR-mediated infection using a dual receptor-specific human H2N2 IAV. Previous genetic screens have defined the host factor landscape of conventional sialic acid-dependent IAV infection (9, 11, 23, 37) and, more recently, identified HLA-DR as the entry receptor for the MHC-II-tropic H18N11 IAV (13). However, these studies used IAV strains restricted to either sialic acid or MHC-II receptor usage, preventing a direct comparison of host factor dependencies between these two receptor-mediated entry pathways. By using a dual receptor-specific IAV in an isogenic, receptor-defined cellular system, our approach enables direct comparison of both infection pathways and receptor-specific host dependencies to be distinguished.

The screen in sialic acid-expressing cells identified a host factor landscape broadly consistent with the established requirements for conventional IAV entry. Enriched candidates included genes involved in sialic acid metabolism and glycosylation, as well as endolysosomal trafficking and acidification, recapitulating key determinants identified in previous genetic screens (11, 23). To our knowledge, this study provides the first genome-wide CRISPR-Cas9 screen using a human H2N2 IAV, the subtype responsible for the 1957 Asian influenza pandemic, providing a resource for exploring subtype-specific determinants of IAV infection. For example, *GALNT6* and *GALNT9*, genes involved in O-linked glycosylation (38), emerged among the enriched candidates. Increasing evidence supports a role for sialylated O-linked glycans in IAV infection (39, 40), with their recognition varying among viral subtypes (39). Although the contribution of O-linked glycosylation to H2N2 infection remains to be established, these candidates illustrate how this dataset could provide a starting point for investigating host processes that may differentially influence infection across IAV subtypes.

The screen performed in HLA-DR-expressing cells identified host dependency factors associated with HLA-DR-mediated IAV infection. Overall, the overlap with the sialic acid screen was limited. At the gene level, this limited overlap could be influenced by the marked difference in the number of hits identified in each screen, likely reflecting the reduced dynamic range of selection in the HLA-DR condition. Notably, genes involved in receptor expression were not shared between the screens, supporting the ability of the approach to capture receptor-associated dependencies. At the pathway level, the sialic acid screen was enriched for processes associated with viral entry, whereas the HLA-DR screen showed a prominent enrichment of pathways related to cellular responses to infection. The recovery of *HLA-DRA* as a top hit nevertheless supported the suitability of the HLA-DR dataset for identifying receptor-associated host factors. To then identify HLA-DR-specific host factors, we integrated both screens into a single HLA-DR-specific score that rewards dependency in the HLA-DR screen while penalizing dependency in the sialic acid screen, and applied guide-support criteria to define a prioritized candidate subset. This integration delivered a focused, ranked list of the host factors most selectively associated with HLA-DR-mediated infection. In contrast, genes shared between the two screens included previously described IAV host factors such as *NXF1*, which is involved in viral mRNA export (41). The recovery of such factors in both screens is consistent with our hypothesis that receptor-specific differences in host requirements are most pronounced during the early stages of infection, whereas post-entry processes are more likely to be shared.

An unexpected feature of the HLA-DR screen was the strong enrichment of multiple components of the IFN-I signaling pathway despite their well-established roles in antiviral immunity (42). Validation experiments showed that disruption of IFNAR2 or STAT1 did not impair either early infection or multicycle IAV replication, indicating that these genes do not function as direct proviral factors during H2N2 infection. Notably, components of the IFN-I signaling pathway have also been recovered in several other survival-based CRISPR-Cas9 screens for IAV (13, 43) and SARS-CoV-2 (44, 45), suggesting that their enrichment represents a recurrent feature of this screening strategy. During IAV infection, infected and bystander cells produce IFN-I, which activate JAK-STAT signaling through IFNAR1 and IFNAR2 to induce expression of IFN-stimulated genes (ISGs) and establish an antiviral state (46). While this response restricts viral spread, it is also accompanied by well-described antiproliferative and cytostatic effects (29). In the context of IAV infection, a recent study has shown that prolonged IFN-I signaling limits epithelial regeneration after IAV infection, delaying tissue repair and increasing susceptibility to secondary bacterial infections (47). Consistent with previous studies, exposure to IFNα and IFNβ markedly inhibited proliferation of DKO-HLA-DR cells, and disruption of IFNAR2 or STAT1 largely abolished this response. Consequently, cells unable to respond to IFN-I acquire a selective growth advantage under the screening conditions, explaining their enrichment in the surviving population. This finding reflects a key feature of survival-based genetic screens: recovered genes include both viral dependency factors and host pathways that influence cellular fitness during infection, highlighting the importance of distinguishing between these when interpreting results from this type of screens.

Among the HLA-DR-specific candidates identified in our comparative analysis, STX2 emerged as the strongest validated host factor for HLA-DR-mediated infection. Using authentic H2N2 virus and a luciferase-expressing H2N2 reporter virus, we consistently observed that depletion of STX2 reduced infection in HLA-DR-expressing cells. Importantly, the same manipulation had little or no effect on infection mediated through sialic acid, indicating that STX2 is not a general IAV host factor but instead contributes selectively to HLA-DR-dependent infection. Likewise, STX2 depletion impaired infection by H2N2-pseudotyped VLPs while leaving VSV-G-mediated entry largely unaffected, further linking its activity to HA-dependent entry rather than to a general defect in endocytosis or reporter virus infection. Our data also provide insights regarding the stage of infection at which STX2 may function. STX2 depletion did not alter surface expression of HLA-DR, excluding its role in modulating receptor abundance under our experimental conditions. Likewise, the total number of cell-associated virions detected shortly after infection remained largely unchanged, suggesting that initial attachment and internalization of virions are not substantially impaired. In contrast, nuclear accumulation of incoming vRNPs was markedly reduced. Together, these observations suggest that STX2 may function during a post-attachment step of HLA-DR-mediated entry, although the precise stage remains to be established.

The known cellular functions of STX2 provide potential mechanistic explanations. STX2 belongs to the syntaxin family of SNARE proteins, which regulate membrane fusion throughout the secretory and endocytic pathways and have been implicated in endosomal recycling and vesicle maturation (31, 33). STX2 could therefore contribute to intracellular trafficking events required for productive HLA-DR-mediated infection. Consistent with this possibility, recent work has shown that STX2 promotes phagosome maturation through fusion with late endosomal and lysosomal compartments (48), suggesting a potential role in regulating vesicle fusion and acidification. Moreover, a recent study has shown that H18 HA-mediated fusion occurs at a lower pH than conventional HAs and requires MHC-II-HA interactions (49). Determining whether MHC-II engagement similarly modulates the fusogenic properties of H2, and whether STX2 contributes to trafficking virion-containing vesicles to compartments that support membrane fusion, may help define the cellular mechanism underlying MHC-II-mediated IAV entry. Further studies will be required to establish the precise contribution of STX2 to this pathway.

Overall, this study provides a framework for dissecting receptor-specific host factor requirements during dual receptor-specific IAV infection. Our findings suggest that MHC-II receptor usage is associated with distinct intracellular host dependencies, exemplified by STX2, supporting the idea that engagement of this alternative receptor may direct incoming virions through intracellular pathways different from those used during conventional sialic acid-mediated entry.

## MATERIALS AND METHODS

### Cell culture

Human lung adenocarcinoma Calu-3 cells (HTB-55), human embryonic kidney 293T (HEK293T) (CRL-3216), and Madin-Darby canine kidney (MDCK) (CCL-34) cells were purchased from the American Type Culture Collection (ATCC).

Calu-3 *HLA-DRA*/*SLC35A1* DKO cells were previously described (16) and reconstituted with either *SLC35A1* (DKO-SLC35A1), *HLA-DRA* plus *HLA-DRB1* (DKO-HLA-DR), or empty vector controls (DKO-Ctrl). HEK293T, MDCK and Calu-3 were maintained in Dulbecco’s modified Eagle medium (DMEM) (41966029, Gibco) supplemented with 10% (v/v) fetal bovine serum (FBS) (A5256701, Gibco) and 100 U/mL penicillin/streptomycin (Pen-Strep) (15070063, Gibco). Calu-3 *HLA-DRA*/*SLC35A1* DKO-derived cell lines were maintained in DMEM containing 20% (v/v) FBS and 100 U/mL Pen-Strep. DKO-SLC35A1 cells were additionally supplemented with 1 mg/mL geneticin (10131035, Gibco), while DKO-HLA-DR and DKO-Ctrl cells were additionally supplemented with 1 mg/mL geneticin and 300 μg/mL hygromycin B (ant-hg-1, InvivoGen). All cells are routinely tested for mycoplasma contamination and confirmed as mycoplasma negative.

### Reconstitution of DKO cells by lentiviral transduction

*SLC35A1* and *HLA-DRA* genes were cloned into the lentiviral vector pLVX-IRES-neomycin (632181, Clontech) via XhoI/BamHI restriction sites, as previously described (13). The pLV-EF1α-HLA-DRB-IRES-hygromycin plasmid was previously described (13). To generate lentiviruses containing either pLVX-SLC35A1-IRES-neomycin, pLVX-HLA-DRA-IRES-neomycin plus pLV-EF1α-HLA-DRB-IRES-hygromycin or corresponding empty vector controls, HEK293T cells were co-transfected with the respective lentiviral plasmids, pMD2.G and pCMVdR8.91 in OptiMEM (31985062, Gibco) using a 3:1 FuGENE® HD Transfection Reagent (E5911, Promega):DNA ratio. At 72 h post-transfection lentivirus-containing supernatants were harvested, centrifuged (3000 r.p.m, 5 min at 4 °C) and filtered through a 0.45 μm filter.

Calu-3 *HLA-DRA*/*SLC35A1* DKO cells were transduced with pLVX-SLC35A1-IRES-neomycin to generate DKO-SLC35A1 cells, with pLVX-HLA-DRA-IRES-neomycin plus pLV-EF1α-HLA-DRB-IRES-hygromycin to generate DKO-HLA-DR cells and with pLVX-IRES-neomycin plus pLV-EF1α-IRES-hygromycin (8513423, Addgene) to generate DKO-Ctrl cells. Cells were inoculated with the respective lentivirus in the presence of 8 μg/mL polybrene. At 24 h post-transduction cells were selected with the corresponding selection antibiotics.

### Viruses

The IAV strains used in this study are A/Japan/305+/1957 (A/JP/57, H2N2) and A/Puerto Rico/8/1934 (A/PR/8, H1N1). The recombinant chimeric H2N2 JP/57 6+2 virus containing the HA and NA segments from A/Japan/305/1957 and the internal segments (PB2, PB1, PA, M, NP, NS) from A/Puerto Rico/8/1934 was previously described (16). Viruses were grown and titered by standard plaque assay on MDCK cells. The recombinant H2N2 (A/JP/57)-*Renilla* reporter virus containing the internal segments (PB2, PB1, PA, M, NP, NS) from A/WSN/1933 (H1N1), the NA segment from A/Japan/305+/1957 and a reporter segment in which the HA-coding sequence is replaced by the *Renilla*-luciferase coding sequence were generated as previously described (50). This virus was grown in MDCK cells expressing the HA from A/Japan/305+/1957, and virus titers were determined by plaque assay in MDCK-H2 (JP/305+/57) cells.

### Biosafety

Experiments performed with wild-type human A/Japan/305+/1957 (H2N2) and recombinant H2N2 JP/57 6+2 carrying the HA and NA segments from A/Japan/305/1957 and the internal segments (PB2, PB1, PA, M, NP, NS) from A/Puerto Rico/8/1934 were performed in a biosafety level 3 laboratory. All other work was carried out at biosafety level 2.

### Protein and glycan detection through flow cytometry

Cells were harvested by trypsinization with 0.25% Trypsin-EDTA (25200-056, Gibco) and stained with LIVE/DEAD Fixable Near-IR Dead Cell Stain (L10119, ThermoFisher) diluted 1:1000 in FACS buffer (PBS, 1 mM EDTA (AM9260G, Invitrogen), 2% BSA) for 30 min at 4 °C. For surface staining, cells were fixed in 3.7% formaldehyde, washed three times with PBS and incubated with primary antibodies or lectins for 1 h at 4 °C. Where applicable, cells were washed three times with PBS and incubated with fluorophore-conjugated secondary antibodies for 30 min at 4 °C. For intracellular staining, cells were fixed, permeabilized with 0.1% w/v saponin for 10 min, and stained with antibodies in FACS buffer for 1 h at 4 °C. Samples were analysed on a BD FACSSymphony flow cytometer, acquiring ≥10,000 events per sample. Data were analysed using FlowJo (v10).

Primary antibodies included anti-human IFNAR2 (21385, PBL Assay Science; diluted 1:100), anti-human HLA-DR (307602, BioLegend; diluted 1:500), and APC-conjugated anti-human HLA-DR (307610, BioLegend; diluted 1:50), Brilliant Violet 421-conjugated anti-human HLA-DR (307635, BioLegend; diluted 1:50). Alexa Fluor 488-conjugated donkey anti-mouse IgG (H+L) was used as secondary antibody diluted 1:1000 when indicated. For α2,3- or α2,6-linked sialic acid detection the biotinylated *Maackia Amurensis II* (MALII) lectin (B-1265, Vector laboratories; diluted 1:100) or the *Sambucus Nigra* (SNA) lectin (B-1305, Vector laboratories; diluted 1:500) followed by secondary staining with APC-labelled streptavidin (405207, Biolegend; diluted 1:500) were used.

### Pseudotyped HIV-based VLP production and infection

Pseudotyped luciferase-encoding HIV-based VLPs were produced by co-transfecting HEK293T cells with 2.8 μg of pNLLuc-AM (51), 0.5 μg of pCAGGS-HA and 0.5 μg of pCAGGS-NA of the indicated IAV strains in OptiMEM supplemented with Viafect (E4981, Promega) in a 1:3 DNA:Viafect ratio. As negative control, HA- and NA-pCAGGS were substituted with 1 ng of pCAGGS-empty vector. Cells were incubated at 37 °C. After 72 h, VLP-containing supernatants were harvested and centrifuged at 3000 r.p.m. for 5 min to remove cell debris. VLP-containing supernatants were treated with 5 μg/ml TPCK-trypsin (T1426, Sigma-Aldrich) for 20 min at 37 °C, followed by treatment with 10 ug/ml soybean trypsin inhibitor (T6522, Sigma-Aldrich) for 20 min at 37 °C. Trypsin-activated VLPs were aliquoted and stored at -80 °C.

Constructs encoding HA and NA from strains A/Japan/305/1957, A/Taiwan/1/1964, A/Brisbane/59/2007 and A/ bat/Peru/33/10 were previously described (13, 16). A/Hong Kong/2286/2017 H3 and N2 (GISAID accession number: HA EPI_ISL_290662) were synthesized as complementary cDNAs (Invitrogen) and cloned into pCAGGS via XhoI/NotI restriction sites.

For VLP infection, cells were seeded in 96-well plates and incubated overnight at 37 °C. The next day, medium was removed and cells were washed once with PBS. Cells were inoculated with 50 μL of undiluted VLPs and incubated 90 min at 37 °C. After that, inoculum was removed, cells were washed once with PBS and 100 μL DMEM supplemented with 10% (v/v) FBS, 100 U/mL Pen-Strep were added. Cells were incubated at 37 °C for 72 h. After that, luciferase signal was measured using ONE-Glo^TM^ Luciferase Assay System (E6110, Promega) according to manufacturer instructions using a Perkin Elmer EnVision device.

### Detection of IAV NP by immunofluorescence staining

Cells were seeded on 96-well plates at 1.5×10^4^ cells/well or on 24-well glass coverslips at 1×10^5^ cells/well and incubated overnight at 37 °C. The next day, cells were washed once with PBS and infected with the indicated IAV strains at the specified MOI in infection PBS (PBS supplemented with 0.3% BSA (A7906, Sigma-Aldrich), 1 mM Ca^2+/^Mg^2+^, and 100 U/mL Pen-Strep) for 1 h at 37 °C. Cells were washed once with PBS, and post-infection DMEM (DMEM supplemented with 0.1% FBS, 0.3% BSA, 20 mM HEPES (H7523, Sigma-Aldrich), and 100 U/mL Pen-Strep) was added. Cells were incubated at 37 °C until the indicated hours post-infection and then washed once with PBS and fixed in 3.7% (v/v) formaldehyde (47608, Sigma-Aldrich) for 20 min at room temperature.

For NP staining, fixed cells were blocked and permeabilized with confocal buffer (PBS containing 50 mM NH_4_Cl (254134, Sigma-Aldrich), 0.1% (w/v) saponin (47036, Sigma-Aldrich), and 2% BSA) for 30 min at room temperature. Cells were washed once with PBS and incubated with mouse monoclonal anti-NP antibody HB-65 (H16-L10-4R5, ATCC) diluted 1:5 in confocal buffer for 1 h at room temperature. Following three PBS washes, cells were incubated with Alexa Fluor 488-conjugated donkey anti-mouse IgG (H+L) highly cross-adsorbed secondary antibody (A-21202, Invitrogen) and DAPI (10236276001, Sigma-Aldrich), both diluted 1:1000 in confocal buffer, for 1 h at room temperature. Cells were washed three times with PBS and one with double-distilled water prior to imaging. Images were acquired using a Leica DMIL microscope and analysed with LAS X (v3.7.4) software.

### Evaluation of virus-induced cytopathic effect

Calu-3 DKO cells were seeded on 96-well plates at approximately 1.5×10^4^ cells/well and incubated 24 h at 37 °C. Cells were washed once with PBS and inoculated with A/Japan/305+/1957 (H2N2) at the indicated MOI in infection PBS for 1 hour at 37 °C. Cells were then washed once with PBS and incubated in post-infection DMEM supplemented with 1 μg/mL TPCK-trypsin (T1426, Sigma-Aldrich) at 37 °C. At 72 h post-infection, cells were washed once with PBS and fixed in 3.7% (v/v) formaldehyde in PBS for 20 min. Fixed cells were washed three times with PBS and stained with crystal violet solution (1% (v/v) crystal violet (61135, Sigma-Aldrich), 10% (v/v) methanol (1.06009, Supelco) in double-distilled water).

### Genome-wide loss of function CRISPR-Cas9 screens

A genome-wide loss-of-function CRISPR-Cas9 screen was performed using the human GeCKOv2 pooled library (1000000048, Addgene) cloned in the lentiCRISPRv2 vector system (21). This library contains 123.411 unique sgRNAs targeting 19.050 human genes, 1.864 microRNAs (miRNAs) and 1.000 non-targeting controls. Lentiviral particles encoding the GeCKOv2 library were produced as previously described (52). Briefly, HEK293T cells were co-transfected with the GeCKOv2 library, psPAX2 (12259, Addgene), and pMD2.G (12260, Addgene) using polyethylenimine (PEI Max) (24765-1, Polysciences). Supernatants were collected 48 h post-transfection, clarified by centrifugation (3000 rpm, 5 min), and filtered through a 0.45 μm filter. Viral titers were determined by transduction as described previously (52). For screening, approximately 1.5×10^8^ DKO-SLC35A1 and DKO-HLA-DR cells were transduced at a MOI of ∼0.3, to achieve ∼300-400× sgRNA coverage throughout the screen. Cells were selected with 1 μg/mL puromycin (ant-pr-1, InvivoGen) for 7 days in DMEM supplemented with 20% (v/v) FBS and 100 U/mL Pen-Strep. Following selection, a fraction of cells (∼1-4×10^8^) was harvested as baseline sample. In parallel, ∼1-2×10^8^ cells were infected with A/Japan/305+/1957 (H2N2) IAV at an MOI of 1. Surviving cells were expanded for approximately 7 days and subsequently collected for genomic DNA extraction. Two independent biological replicates were performed per cell line. Due to reduced sgRNA enrichment observed in the first DKO-HLA-DR replicate, a modified infection strategy was applied in the second replicate, consisting of two sequential infections at MOI 5, with re-infection at 96 h post-infection. Genomic DNA was isolated using the Quick-DNA Midiprep Plus Kit (D4075, Zymo Research) according to the manufacturer’s instructions. sgRNA sequences were amplified by PCR (26 cycles) as previously described (52) using NEBNext High-Fidelity PCR Master Mix (M0541S, New England Biolabs), incorporating Illumina adapters and sample barcodes. Primers used for sgRNA amplification and sequencing library preparation are described in Supp. Table S6. PCR products (∼270 bp) were purified by agarose gel electrophoresis and gel extraction. Libraries were pooled, quantified, and sequenced on an Illumina NextSeq 2000 platform (single-end 100 bp reads) with 20% PhiX spike-in at the Functional Genomics Center Zurich. Sequencing data were analysed using the PinAPL-Py web tool (22), applying size-factor normalization and the αRRA algorithm for gene-level ranking.

### Generation of HLA-DR-specific gene ranking

To integrate the gene-level results of the screens in DKO-SLC35A1 and DKO-HLA-DR cells and nominate candidates for experimental validation, we developed the HLA-DR-specific score. This metric combines the statistical significance of gene enrichment with the number of significantly enriched sgRNAs identified in each screen. Genes were included in the ranking if their αRRA value in the DKO-HLA-DR screen was <1.0 and they had at least one significantly enriched sgRNA in that screen. sgRNA enrichment significance was determined using Benjamini–Hochberg correction at an FDR threshold of 0.01. The HLA-DR component of the score rewarded genes displaying strong enrichment and support from multiple significant sgRNAs, whereas the sialic acid component was incorporated as a penalty to reduce the ranking of genes also enriched in the DKO-SLC35A1 screen. The final HLA-DR-specific score was calculated as: HLA-DR strength [-log_10_(αRRA_DKO-HLA-DR_ + ε) x sqrt (n sig. sgRNA_DKO-HLA-DR_ / 2)] – Sialic acid penalty [-log_10_(αRRA_DKO-SLC35A1_ + ε) x ((n sig. sgRNA_DKO-SLC35A1_ + 1) / 2)], where α*RRA* represents the robust rank aggregation score generated by PinAPL-Py, *n sig. sgRNA* denotes the number of significantly enriched sgRNAs for each gene in the respective screen, and ε = 10^-10^ was added to avoid undefined logarithms. For score calculation and eligibility filtering, missing αRRA values were replaced with 1.0 and missing significant-sgRNA counts with 0. Genes were subsequently ranked by decreasing HLA-DR-specific score. To define a prioritized candidate subset, genes were required to satisfy two guide-support criteria: (i) at least two significantly enriched sgRNAs in the DKO-HLA-DR screen and (ii) no more than one significantly enriched sgRNA in the DKO-SLC35A1 screen.

### Generation of polyclonal KO cell lines

Polyclonal KO cell populations were generated by lentiviral transduction of lentiCRISPRv2 vectors encoding sgRNAs targeting the indicated genes, as previously described (52). Briefly, complementary sgRNA oligonucleotides were annealed, phosphorylated, and ligated into BsmBI-digested lentiCRISPRv2 vector (Addgene #52961) according to (21). Correct sgRNA insertion was confirmed by Sanger sequencing. All sgRNA sequences are listed in Supp. Table S7. Where indicated, three independent sgRNAs per gene were pooled to increase KO efficiency and reduce sgRNA-specific effects. Lentiviral particles were produced by co-transfecting HEK293T cells with psPAX2, pMD2.G, and an equimolar mixture of the three corresponding lentiCRISPRv2-sgRNA constructs. Target cells were transduced for 24 h and selected with puromycin (1 μg/mL) for 7 days to generate polyclonal KO cell populations. Following selection, cells were maintained for no more than five passages before downstream analyses.

### Measurement of virus growth kinetics

Cells were seeded in 96-well plates at 1.5×10^4^ cells/well and incubated overnight at 37 °C. For infection, cells were washed once with PBS, inoculated with the indicated virus strain at the specified MOI in infection PBS and incubated for 1 h at 37 °C. Inoculum was removed, cells were washed once with PBS and post-infection DMEM supplemented with 1 μg/mL TPCK-trypsin was added. At the indicated times post-infection, virus titers were determined by standard plaque assay.

### Cell growth measurement

Cells were seeded on 96-well plates at 1.2×10^4^ cells/well and incubated 24 h at 37 °C. The next day, cells were stimulated with the corresponding concentration of IFNα2 (NPB2-34971; Novus Biologicals) or IFNβ (11420-1, PBL Assay Science) diluted in DMEM supplemented with 10% v/v FBS and 100 U/mL Pen-Strep. To monitor cell growth, cell confluency was measured every 8 h in an Incucyte S3 device for four days. Cell confluency was estimated using the Incucyte S3 software.

### IAV M segment detection through digital PCR

Cells were seeded in 24-well plates at 1×10^5^ cells/well and incubated at 37 °C. After 24 h, cells were washed once with PBS and infected with the respective virus at the indicated MOI in infection PBS for 1 h at 37 °C. Cells were washed once with PBS, post-infection DMEM was added, and cells were incubated at 37 °C. After 6 h, cells were lysed and RNA was extracted using the ReliaPrep™ RNA Cell Miniprep System (Z6010, Promega) according to manufacturer’s instructions.

Detection and quantification of IAV M segment was performed using the QIAcuity Digital PCR System and QIAcuity OneStep Advanced Probe Kit (250132, Qiagen) as previously described (53). The following primers and probes were used: forward primer 5’-TGG AAT GGC TAA AGA CAA GAC CAA T-3’; reverse primer 5’-AAA GCG TCT ACG CTG CAG TCC-3’; probe TTT GTK TTC ACG CTC ACC GTG CCC. Absolute RNA concentration (copies/µL) for each sample was calculated using the QIAcuity Software Suite.

### IAV-*Renilla* reporter virus infection

Cells were seeded on 96-well white plates (CLS3917, Corning) at 1.5×10^4^ cells/well and incubated overnight at 37 °C. Cells were then washed once with PBS and inoculated with the IAV-*Renilla* reporter virus at the indicated MOI in infection PBS for 1 h at 37 °C. After that, cells were washed once with PBS and post-infection DMEM supplemented with 6 µM Enduren^TM^ Live Cell Substrate (E6481, Promega) was added. Luciferase signal was measured every hour with a Perkin Elmer EnVision device for 8 h.

### siRNA-mediated gene silencing

For siRNA transfection, cells were reverse transfected with 30 nM of the corresponding siRNA (1027415, Qiagen) diluted in OptiMEM (31985062, Gibco) using Lipofectamine RNAiMax (13778150, Invitrogen), following manufacturer’s instructions. The sequences of the siRNAs used in this study can be found in Supp. Table S8. Downstream assays with siRNA-transfected samples were performed at 48 h post-transfection.

### Cell viability assay

Cell viability of siRNA-treated samples was measured at 48 h post-transfection using the CellTiter-Glo Luminescent Cell Viability Assay kit (G7571, Promega), following the manufacturer’s guidelines. Luminescent signal was measured in a Perkin Elmer EnVision device.

### Detection of STX2 by confocal microscopy

Cells were seeded on 24-well glass coverslips at approximately 1×10^5^ cells/well. Cells were washed once with PBS and fixed with 4% paraformaldehyde (15710, Electron Microscopy Science) for 10 min at room temperature. After that, cells were washed three times with PBS and permeabilized in confocal buffer for 1 h at room temperature. Cells were washed once with PBS and incubated with the rabbit polyclonal anti-human syntaxin-2 purified antibody (110123, Synaptic Systems) diluted 1:100 in confocal buffer for 1 h at room temperature. Cells were washed three times with PBS and incubated with the Alexa Fluor 647-conjugated donkey anti-rabbit IgG (H+L) highly cross-adsorbed secondary antibody (A-31573, Invitrogen) and DAPI, both diluted 1:1000 in confocal buffer, for 1 h at room temperature. Cells were washed three times with PBS, rinsed with double-distilled water and mounted on a microscope glass slide in ProLong™ Gold Antifade Mountant (P36930, Invitrogen). Images were acquired in a Leica SP8 confocal laser scanning microscope. Confocal image analysis was performed using Imaris v11.0.0 software. STX2-positive structures were identified by surface rendering using identical thresholding parameters across all conditions. Total STX2 fluorescence was quantified as the Intensity Sum of all detected surfaces per field of view. As similar cell densities were present across imaging fields, no normalization to cell number was performed.

### Pseudotyped VSVΔG VLPs production and infection

Pseudotyped luciferase-encoding VSVΔG VLPs were produced by co-transfecting HEK293T cells with pCAGGS-H2 and pCAGGS-N2 plasmids (0.5 μg each) encoding the HA and NA proteins of A/Japan/305/1957 (H2N2) or with pCAGGS-VSV-G (1 μg) using ViaFect Transfection Reagent (E4981, Promega) at a DNA:ViaFect ratio of 1:3 in OptiMEM. Cells transfected with pCAGGS empty vector served as a negative control. At 24 h post-transfection cells were infected with a recombinant VSVΔG/G*-luciferase (EH1020-PM, Kerafast) at an MOI of 4 for 1 h at 37 °C. Inoculum was removed and cells were incubated in DMEM supplemented with 1 µg/mL anti-VSV-G antibody (EB0010, Kerafast) at 37 °C to neutralize residual input virus. After 48 h, VLP-containing supernatants were harvested and centrifuged at 3000 r.p.m. for 5 min to remove cell debris. VLPs pseudotyped with IAV HA glycoproteins were activated with 5 μg/ml TPCK-trypsin for 20 min at 37 °C, followed by treatment with 10 μg/ml soybean trypsin inhibitor (T6522, Sigma-Aldrich) for 20 min at 37 °C. VLPs were aliquoted and stored at -80 °C.

For infection assays, 1.5×10^4^ cells/well were seeded in 96-well plates and cultured overnight at 37 °C. The next day, medium was removed and cells were washed once with PBS. Cells were inoculated with 50 μL undiluted VLPs and incubated 90 min at 37 °C. Inoculum was removed and cells were washed once with PBS and incubated in DMEM supplemented with 10% v/v FBS, 100 U/mL Pen-Strep 37 °C. After 7 h, luciferase activity was measured using ONE-Glo^TM^ Luciferase Assay System (E6110, Promega) according to manufacturer’s instructions using a Perkin Elmer EnVision device.

### Quantification of IAV NP through confocal microscopy

Cells were seeded onto glass coverslips in 24-well at 1×10^5^ cells/well and incubated overnight at 37 °C. Cells were washed once with PBS and infected with recombinant H2N2 JP/57 6+2 virus at MOI of 40 in infection PBS supplemented with either 100 μg/ml cycloheximide (C4859, Sigma-Aldrich), 10 nM bafilomycin A1 (B1793, Sigma-Aldrich) or the corresponding concentration of DMSO (D2650, Sigma-Aldrich) for 1 h at 37 °C. Following inoculum removal, cells were incubated in post-infection DMEM containing the respective compounds. At 4 h post-infection, cells were fixed with 4% paraformaldehyde for 20 min at room temperature.

Fixed cells were permeabilized in confocal buffer for 1 h at room temperature and incubated with mouse monoclonal anti-NP antibody HB-65 diluted 1:10 in confocal buffer for 1 h at room temperature. Following three PBS washes, cells were incubated with Alexa Fluor 488-conjugated donkey anti-mouse IgG (H+L) highly cross-adsorbed secondary antibody (A-21202, Invitrogen) and DAPI (10236276001, Sigma-Aldrich), both diluted 1:1000 in confocal buffer, for 1 h at room temperature. Cells were washed three times with PBS, rinsed with double-distilled water and mounted on a microscope glass slide in ProLong™ Gold Antifade Mountant. Coverslips were imaged in a Leica SP8 confocal laser scanning microscope. Confocal image analysis was performed in Imaris v11.0.0 software. NP-positive puncta were identified and quantified by spot detection (estimated diameter, 0.3 μm; quality threshold, > 2.0) and nuclei were segmented from the DAPI signal using surface detection (surface grain size, 0.5 μm; manual threshold, 24). NP puncta were subsequently classified as intra-or extranuclear according to their localization relative to the segmented nuclear surfaces.

## ACKNOWLEDGEMENTS

We thank the Functional Genomics center of Zurich for the amplicon sequencing of the CRISPR-Cas9 screen samples. This study was funded by the Swiss National Science Foundation (grant number 310030_204166 to SSt) and the UZH Candoc Grant (grant number FK-25-019 to LMAF).

## AUTHOR CONTRIBUTION

L.M.A.-F. and S.S. designed the experiments. L.M.A.-F., U.K., and I.G. performed the experiments. L.P. designed and performed bioinformatic analysis. P.M.B. generated reagents for the study. L.M.A.-F. and S.S. wrote the manuscript.

## DATA AVAILABILITY

All data needed to evaluate the conclusions in the paper are present in the paper and/or the Supplementary Materials.

---

**Supplementary Figure 1.**
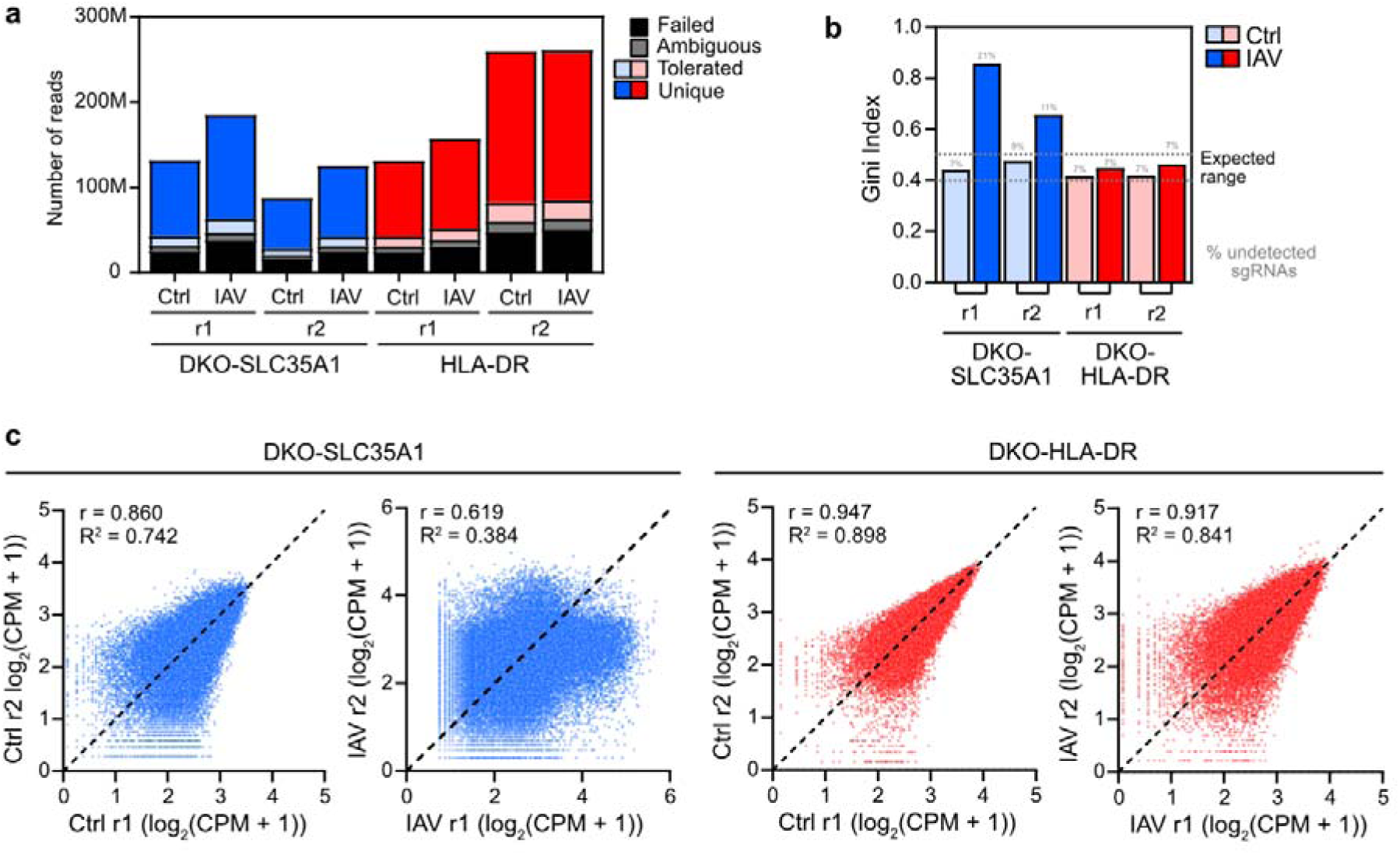
Quality control metrics for genome-wide CRISPR-Cas9 screening. **(a)** Sequencing read counts and mapping statistics for each screen replicate and experimental condition. Bar plots show the total number of sequencing reads classified as *unique*, *ambiguous*, *tolerated*, or *failed* based on alignment to the sgRNA library reference. **(b)** Gini indexes were calculated from sgRNA read count distributions for each condition and biological replicate as a measure of library representation. The percentage of undetected sgRNAs (zero read counts) is indicated for each sample. **(c)** Reproducibility between biological replicates. Scatter plots show sgRNA abundance expressed as log_₂_ (CPM + 1) for replicates 1 and two in pre-selection (Ctrl) and post-selection (IAV) populations. Pearson correlation coefficient (*r*) and the corresponding coefficient of determination (*R*²) are indicated for each comparison.

**Supplementary Figure 2.**
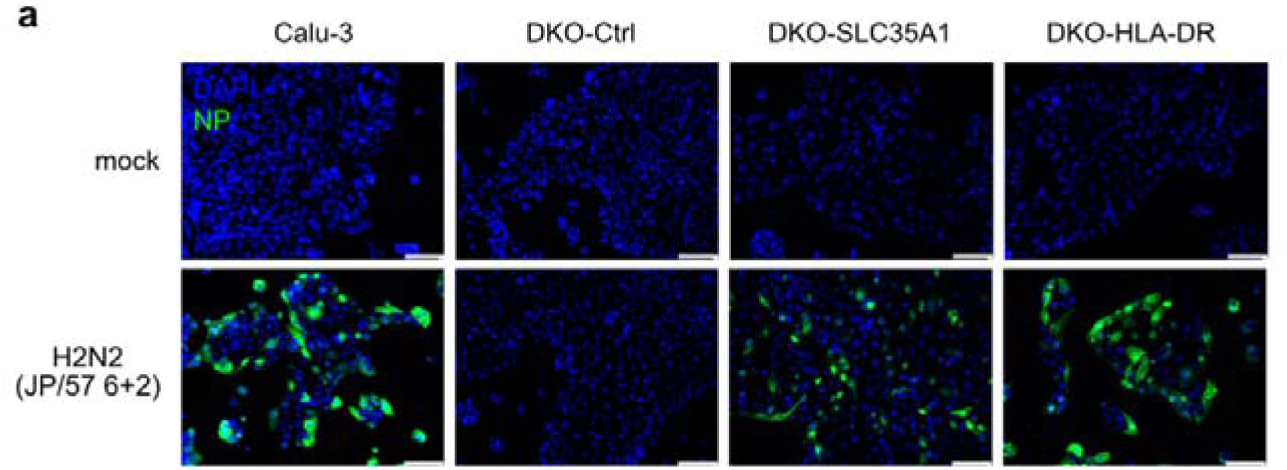
Validation of receptor-specificity for the recombinant H2N2 JP/57 6+2 virus. **(a)** Calu-3 and Calu-3 *HLA-DRA*/*SLC35A1* DKO-derived cell lines were infected with H2N2 JP/57 6+2 virus (MOI 5) for 7 h. Viral nucleoprotein (NP, green) and nuclei (DAPI, blue) were detected by immunofluorescence.

**Supplementary Figure 3.**
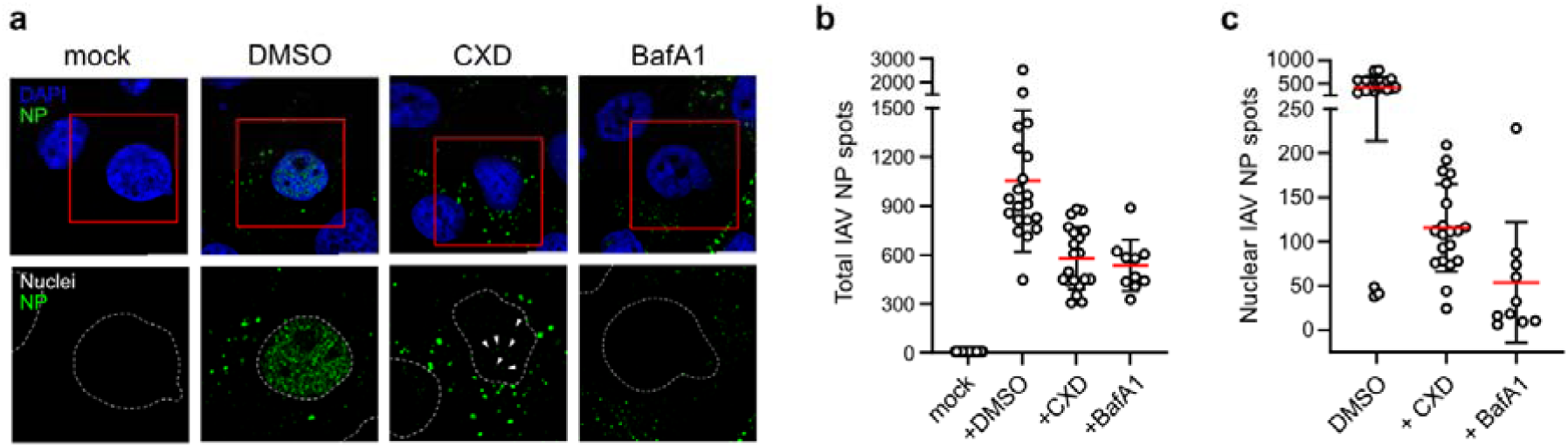
Effect of bafilomycin A1 on IAV entry. **(a)** As described for fig. 5, siRNA-transfected cells were infected with PR8-JP57 (6+2) at MOI 40 in the presence of cycloheximide (100 µg/mL), bafilomycin A1 (10 nM) or DMSO. At 4 h post-infection, cells were fixed, permeabilized and stained for viral NP (NP; green) and nuclei (DAPI; blue or dashed while line). Representative confocal images from n = 2 (n = 1 for the bafilomycin A1-treated sample) independent experiments are shown. Scale bar = 10 µm. **(b)** NP-positive puncta were identified and quantified by spot detection in Imaris. **(c)** For intranuclear NP quantification, nuclei were segmented based on the DAPI signal and NP puncta inside the nuclear surface were quantified. (b) Each datapoint represents one confocal image from n = 2 (mock, DMSO and CXD) or n = 1 (BafA1) independent experiments. Means (red) ± s.d. is shown.

